# Reading negative action verbs: one or two-step processing within the primary motor cortex?

**DOI:** 10.1101/2022.10.25.513652

**Authors:** W Dupont, C Papaxanthis, L Lurquin, F Lebon, C Madden-Lombardi

## Abstract

Controversy persists regarding the representation of negated actions, specifically concerning activation and inhibitory mechanisms in the motor system, and whether this occurs in one or two steps. We conducted two experiments probing corticospinal excitability (CSE) and short-interval intracortical inhibition (SICI) in the primary motor cortex at different latencies while reading affirmative and negative action sentences.

Twenty-six participants read action and non-action sentences in affirmative or negative forms. Using transcranial magnetic stimulation, we probed CSE in hand muscles at rest and at several latencies after verb presentation. We observed a greater CSE for action sentences compared to non-action sentences, regardless of verb form.

In experiment two, nineteen participants read affirmative and negative action sentences. We measured CSE and SICI at short and long latencies after verb presentation. CSE was greater for affirmative and negative action sentences at both latencies compared to rest. SICI did not change at the short latency but increased at longer latencies, regardless of verb form.

Our results lend partial support for a two-step model, as negated actions showed the same motor excitability as affirmed actions with no additional inhibition at early latencies. Later neural differences between affirmative and negative actions may occur outside the primary motor cortex.

**Significant statement:** In two TMS experiments, we probed corticospinal excitability and short-interval intracortical inhibition in the primary motor cortex at different latencies while subjects read affirmative and negative action sentences. Consistent with an embodied view of language comprehension, our results demonstrate that reading about actions indeed activates the motor system, and this for both negative and affirmative sentences. Our results lend partial support for a two-step model of negation, as negated actions showed the same increase in motor excitability as affirmed actions, with no additional inhibition at early latencies. This suggests that the motor system contributes to comprehension by simulating the negated or affirmed action. Later neural differences between affirmative and negative actions may occur outside the primary motor cortex.

## Introduction

Over the past few decades, a growing body of research has established a clear link between perceptual-motor and language processes (Barsalou 1999, 2008; Barsalou et al. 2003; Gallese and Lakoff 2005; Decety and Grèzes 2006; Zwaan and Taylor 2006; Fischer and Zwaan 2008; Jeannerod 2008; Glenberg and Gallese 2012). Such research suggests that language, along with other cognitive functions, is processed in a distributed manner that includes areas of the brain that were thought to be reserved for perception and action. Corroborating this recent distributed view of language processing, the involvement of the motor system during the comprehension of action language has been reported by a variety of methodologies, such as fRMI (Hauk et al. 2004; Van Dam et al. 2010), EEG (Pulvermüller et al. 2001; Kellenbach et al. 2002; Xu et al. 2016; Beres 2017), psychophysics (Pulvermüller et al. 2001; Rabahi et al. 2012, 2013; Andres et al. 2015; Klepp et al. 2019), and Transcranial Magnetic Stimulation (TMS) (Papeo et al. 2009, 2015; Labruna et al. 2011; Innocenti et al. 2014).

While it is clear that action language engages the motor system, this involvement seems to depend on various experimental elements, such as temporal factors (Borreggine and Kaschak 2006; Kaschak and Borreggine 2008), choice of stimuli (Papeo et al. 2009), and contextual factors (Raposo et al. 2009; van Dam et al. 2010; Zwaan et al. 2010; Van Dam et al. 2012, 2014; Gilead et al. 2013; Beauprez et al. 2018). One contextual factor that may modulate the activation of the motor system during action reading is negation. Negation is a linguistic operator used to reject, deny or express non-existence. Processing negation seems to require additional resources, as negated sentences yield longer processing times and higher error rates than affirmative sentences (Clark 1974; Carpenter and Just 1975; Lüdtke 2006; Kaup, Zwaan, et al. 2007). Traditional (amodal, non-distributed) theories of language comprehension could easily explain these results in terms of the negation operator as an extra-linguistic processing step. Distributed theories of language comprehension have also offered explanations for negation processing in terms of a simulation of the negated situation. Some researchers posit that this can be done in a single step (Mayo et al. 2004; Nieuwland and Kuperberg 2008), whereas others postulate a two-step process to account for the negated information (Kaup, Yaxley, et al. 2007; Lüdtke et al. 2008).

According to one-step models, the negated action representation is immediately obstructed, as reflected in decreased behavioral performance and reduced activation of the primary motor and premotor cortices for negative compared to affirmative sentences (Carpenter et al. 1999; Tettamanti et al. 2008; Christensen 2009; Tomasino et al. 2010; Aravena et al. 2012; Bartoli et al. 2013; Foroni and Semin 2013; García-Marco et al. 2019). Two-step models stipulate that both affirmative and negative representations are engaged after reading negative sentences. Specifically, the representation of the action to be negated is engaged in the first step (as in affirmed actions), whereas the representation of the actual situation with the negated action is engaged in a second step. For instance, “not smoking” would first activate a simulation of smoking, and then a simulation without smoking. The initial representation of the to-be-negated information allows comprehension of what has been negated. Without this step, it would be difficult to comprehend the difference between, for example, “not smoking” and “not swimming”. This idea is based on studies reporting initial faster reaction times for the negated content, followed later by faster reaction times for the actual situation described in the sentence (Kaup et al. 2005, 2006; Kaup, Yaxley, et al. 2007; Lüdtke et al. 2008; Anderson et al. 2010; Scappini and Delfitto 2015).

The high temporal resolution of TMS provides a relevant tool to probe the involvement of the motor system in language processing, and it may help to shed light on the neural mechanisms underlying negation. TMS studies showed that corticospinal excitability, a marker of the motor system’s activation, increases during affirmative action verb reading (Papeo et al. 2009; Labruna et al. 2011; Innocenti et al. 2014). Furthermore, Papeo et al. (2016) reported that action verbs presented in the affirmative form elicited greater corticospinal excitability than their negative counterparts at 250, 400, and 550 milliseconds after presentation. These findings suggest early inhibitory mechanisms during the reading of negative action sentences, extending previous EEG studies (De Vega et al. 2016; Beltrán et al. 2019, 2021; Liu et al. 2020).

While the literature on negation processing suggests that inhibitory mechanisms modulate the motor system’s involvement, it remains unclear whether this occurs in one or two steps within the primary motor cortex (M1), and how facilitatory and inhibitory mechanisms might interact. In the current study, we performed two TMS experiments to investigate the neural modulation along the corticospinal track and within M1 during the reading of affirmative and negative action sentences. In a first experiment, we measured MEP amplitude, by means of single-pulse TMS, at different latencies after verb presentation to determine whether the corticospinal excitability increased or/and temporally shifted for negated compared to affirmed action sentences. In a second experiment, we probed the modulation of corticospinal excitability and short-interval intracortical inhibition (SICI), by means of single and paired-pulse TMS respectively, delivered at short and long latencies after verb presentation. In the case of one-step processing, we would expect an immediate inhibition for negated sentences compared to affirmed sentences, i.e., greater SICI. In the case of two-step processing, we would expect the negated sentences to show an increase of corticospinal excitability in comparison to rest at early latencies, followed by an inhibition only at longer latencies.

## Material and method

### Experiment 1

#### Participants

Twenty-six healthy right-handed individuals (13 women; mean age = 22.8 years-old; range 18-28) participated in experiment 1. Participants’ handedness was assessed by the Edinburgh inventory (range 0.25-1; Oldfield 1971). All participants were French native speakers and had normal or corrected-to-normal vision, without neurological, physical or psychiatric pathology. The experimental protocol and procedures were in accordance with the Declaration of Helsinki and were granted ethics committee approval (excluding pre-registration; CPP SOOM III, ClinicalTrials.gov Identifier: NCT03334526).

#### Stimuli

One hundred and sixty (n=160) French sentences were generated; half (n=80) referring to hand actions (e.g., “The potato is cooked, I peel it”) and half (n=80) describing non-actions (e.g., “The potato is cooked, I leave it”) or actions not implicating the hand (e.g., “I kick it”), corresponding to the experimental conditions Action and Non-Action, respectively. All sentences were presented in the first-person present tense and were created so that the target verb appeared at the end of the sentence. The final pronoun-verb segment was presented alone on a subsequent screen (Figure 1). The target verbs were presented either in affirmative form (n=40) (e.g., “The potato is cooked, I peel it”) or in negative form (n=40) (e.g., “The potato is cooked, I don’t peel it”), corresponding to the experimental conditions Affirmative and Negative form, respectively. All stimuli order was randomized and counterbalanced.

**Figure 1.**
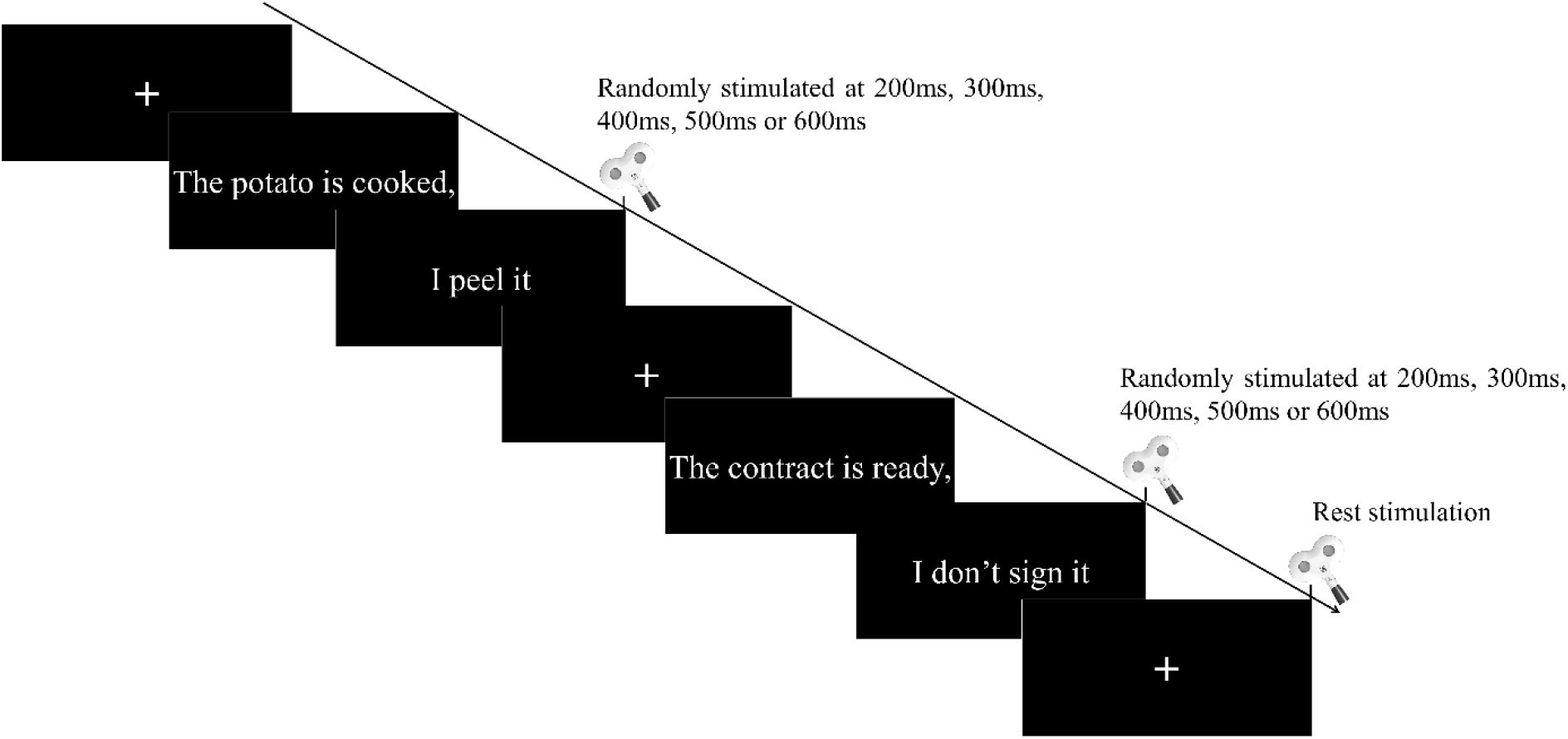
Experimental procedure showing an affirmative, a negative and a rest trial. Sentences were presented in a Negative or Affirmative form. Single-pulse TMS was triggered over the left hemisphere at rest or at various delays after the verb presentation, yielding motor-evoked potentials that were recorded in the right index finger.

Various psycholinguistic factors (e.g., written frequency, number of letters and syllables, spelling neighbors and semantic analysis) were controlled for the verb conditions using the Lexique.org database (New et al., 2004). The statistical tests carried out showed no significant difference between the conditions for all the psycholinguistic factors (See Supplementary section).

#### Procedure

Participants sat in an armchair and read the sentences on a 19-inch LCD monitor. A custom-made software synchronized the presentation of the stimuli, the TMS pulses, and the electromyographic (EMG) recording. Participants were instructed to stay still while they silently read the sentences. Corticospinal excitability was assessed at rest and during the reading tasks at various stimulation latencies (200, 300, 400, 500 or 600 ms after the verb onset). These latencies were chosen based on previous electroencephalogram and TMS studies showing 200ms to 400ms as latencies for action semantic processing in parietal and frontal regions (Pulvermüller et al. 2001; Kellenbach et al. 2002; Hauk et al. 2006; Beres 2017). The latest latencies (500 and 600ms) were added to catch a potential late shift in the activation latency for negative compared to affirmative sentences.

A familiarization session was conducted before the experimental session, in which participants saw four trials, each starting with a fixation cross (500ms), followed by a sentence (3000ms), the pronoun-verb target (2000ms), then a break before the next trial (5000ms). Once the participant understood this procedure, four new practice trials were presented, now including TMS pulses after the appearance of the target verb.

The experimental session was divided into 8 blocks of 43 trials each, yielding 344 trials in total during the experiment. Among the 43 trials in each block, 3 trials included only TMS pulses at rest (upon presentation of the fixation cross), which served as reference stimulations and allowed for comparisons across experimental conditions in that block. For the 40 remaining trials, the TMS pulses were delivered after the verb presentation. In each block, the number of stimulations was equally distributed across all conditions (Action vs. Non-action and Negative vs. Affirmative), and also across the five different stimulation latencies. Moreover, to avoid mental fatigue and lack of concentration, we introduced a two-minute break between each block.

#### EMG recording and TMS set-up

The EMG signal was recorded using 10mm-diameter surface electrodes (Contrôle Graphique Médical, Brice Comte-Robert, France) placed over the right first dorsal interosseous (FDI) muscle. Before placing the electrodes, the skin was shaved and cleaned to reduce EMG signal noise (<20μV). EMG signals were amplified and bandpass filtered on-line (10-1000 Hz, Biopac Systems Inc.) and digitized at 2000 Hz for off-line analysis. We calculated the EMGrms for further analysis. Single-pulse TMS was delivered with a figure-eight coil (70 mm in diameter) connected to a Magstim 200 stimulator (Magstim Company Ltd, Whitland). The coil was positioned over the motor area controlling the right FDI muscle. The coil rested tangential to the scalp with the handle pointing backward and laterally at a 45° angle from the midline. Mappings of each participant’s motor cortex determined the optimal coil position (hotspot), where stimulation evoked the highest and most consistent MEP amplitude for the FDI muscle. We measured the resting motor threshold, which is the minimal TMS intensity to induce a peak-to-peak MEP amplitude of 50 μV in the right FDI muscle for 5 trials out of 10 (Rossini et al. 2015). During the experimental session, the TMS intensity was set at 130% of individual resting motor threshold.

#### Data and statistical analysis

The EMG signal was extracted with Matlab (The MathWorks, Natick, Massachusetts, USA) and the peak-to-peak MEP amplitude was measured for all conditions. Before analysis, we removed data falling 2 standard deviations (SDs) above or below individual means for each condition (4.40% of all trials). Then, the individual mean MEP amplitude for each condition was normalized to rest (% to rest). Two participants were excluded due to extreme values (i.e., over 2 SDs from the group mean), leaving 24 participants in the final analysis. Statistics and data analyses were performed using the Statistica software (Stat Soft, France).

Data normality and sphericity were verified using Shapiro-Wilk and Mauchly tests. In order to assess a general shift in the increase of corticospinal excitability during negated action processing, we first performed a repeated measures ANOVA with Action (Action vs. Non-Action), Polarity (Affirmative vs. Negative) and Latency (200, 300, 400, 500, vs. 600ms) as within-subject factors. The semantically-driven increase in corticospinal excitability that we were interested in measuring would likely be ephemeral, and therefore not present at all latencies. As this increase would likely vary across conditions and participants, we then selected the latency for which the MEP amplitude was at its peak. More specifically, we isolated the highest increase in MEP amplitude among the five stimulation latencies for each participant in each condition. This subject-specific peak method accommodates individual variability in the latency of action semantic processing, and affords probing the motor system when the action representation is active. We submitted these peaks to a repeated measures ANOVA with Action (Action vs. Non-Action) and Polarity (Affirmative vs. Negative) as within-subject factors. Then, one-sample t-tests were used to compare condition MEPs to zero (rest). Finally, to be certain that our results were not biased by muscular activity preceding the TMS pulse, a Friedman ANOVA was used to compare the EMGrms (100ms-window prior the artifact) between rest and our experimental conditions (See Supplementary section). The data are presented as mean values (±standard deviation) and the alpha value was set at 0.05.

### Experiment 2

#### Participants

Nineteen healthy right-handed individuals (7 women; mean age = 22.05 years-old; range 18-27) participated in the experiment. Participants’ handedness was assessed by the Edinburgh inventory (range 0.2-1; Oldfield 1971). All participants were French native speakers with normal or corrected-to-normal vision, without neurological, physical or psychiatric pathology.

The experimental protocol and procedures were in accordance with the Declaration of Helsinki and were granted ethics committee approval (excluding pre-registration; CPP SOOM III, ClinicalTrials.gov Identifier: NCT03334526).

#### Procedure and stimuli

Stimuli and task were identical to Experiment 1, except for two features. First, we only presented affirmative and negative action sentences (no non-action sentences). Second, we assessed corticospinal excitability and short-interval intracortical inhibition (SICI) at rest or at short and long latencies after verb presentation. For the short latencies, we determined which of 3 latencies (200, 300 and 400 ms) yielded the highest MEP amplitude (single-pulse TMS) on a set of practice trials at the beginning of the experiment for each participant, and we used this peak latency as their short latency to probe CSE and SICI during the experiment. For the long latency, we stimulated at 700, 800 and 900ms after verb presentation for single- and paired-pulse TMS during the experiment, then chose the latency that induced the greatest increase of corticospinal excitability (single-pulse TMS) and the greatest SICI (paired-pulse TMS) for each individual and each condition for analyses.

Twenty single and twenty paired TMS pulses were delivered per condition (rest, affirmative verb, negative verb) at 4 latencies: one individualized short latency, and 3 long latencies (700, 800 and 900 ms).

#### EMG recordings and TMS set-up

The EMG recordings and TMS set-up were almost identical to Experiment 1, except for the TMS parameters and measurements described just above. TMS responses were measured via an adaptive threshold-tracking technique. We used an available online freeware (TMS Motor Threshold Assessment Tool, MTAT 2.0) based on a maximum-likelihood Parameter Estimation by Sequential Testing. We first determined the individual MepTarget amplitude, which is half of the individual maximum MEP amplitude at rest. We set the conditioning stimulation (CS) at 60% of resting motor threshold. Then, we modulated the TMS stimulation intensity of the test stimulation (TS) to match the MepTarget amplitude for each condition. Therefore, we did not analyze the MEP amplitude as in Experiment 1, but rather the intensity of the TS stimulation, namely the percentage of Maximal Stimulator Output (%MSO). This approach avoids the recurrent problem of a floor effect in SICI protocols, where inhibitory networks are excessively powerful and the MEPs are occluded, preventing any analysis (Cirillo and Byblow 2016; Cirillo et al. 2018).

#### Data and statistical analysis

Statistics and data analyses were performed using the Statistica software (Stat Soft, France). For corticospinal excitability, we normalized the single-pulse TS intensity of each experimental condition to the single-pulse TS intensity at rest (%). The amount of SICI (expressed in INH%) was quantified for each condition using the following equation (Fisher et al., 2002):

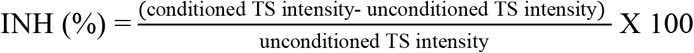

This ratio for the experimental condition was then normalized to the ratio at rest (Δ). Data normality and sphericity were verified using Shapiro-Wilk and Mauchly tests. As in the first experiment, we selected the timing for which the average corticospinal excitability was highest for an individual, and compared MEPs for each condition at zero (rest) using one-sample t-tests. Then, to assess the influence of Latency and Polarity on corticospinal excitability, we realized a 2 by 2 repeated-measures ANOVA with Latency (Short vs. Long) and Polarity (Affirmative vs. Negative) as within-subject factors. Using the same peak latency that was selected in the corticospinal excitability measure, we performed the same statistical design on SICI. Finally, to be certain that our results were not biased by muscular activity preceding the TMS pulse, we compared the EMGrms (100ms-window prior to TMS artifact) between rest and our experimental conditions with Friedman ANOVA (See Supplementary section). The data are presented as mean values (± standard deviation) and the alpha value was set at 0.05.

## Results

### Experiment 1

We observed a main effect of Latency (F_4,92_=9.928, p<0.000, ηp^2^=0.301) and an interaction between Action and Latency (F_4,92_=2.984, p=0.022, ηp^2^=0.114), with greater corticospinal excitability for action compared to non-action sentences only at 300ms (p=0.009). However, we did not observe a main effect of Action (F_1,23_=1.545, p=0.226, ηp^2^=0.062) or Polarity (F_1,23_=0.034, p=0.564, ηp^2^=0.014), and there were no interactions between Polarity and Latency (F_4,92_=0.650 p=0.628, ηp^2^=0.027) nor among Action, Polarity and Latency (F_4,92_=0.391 p=0.814, ηp^2^=0.016) (See Figure 2).

**Figure 2:**
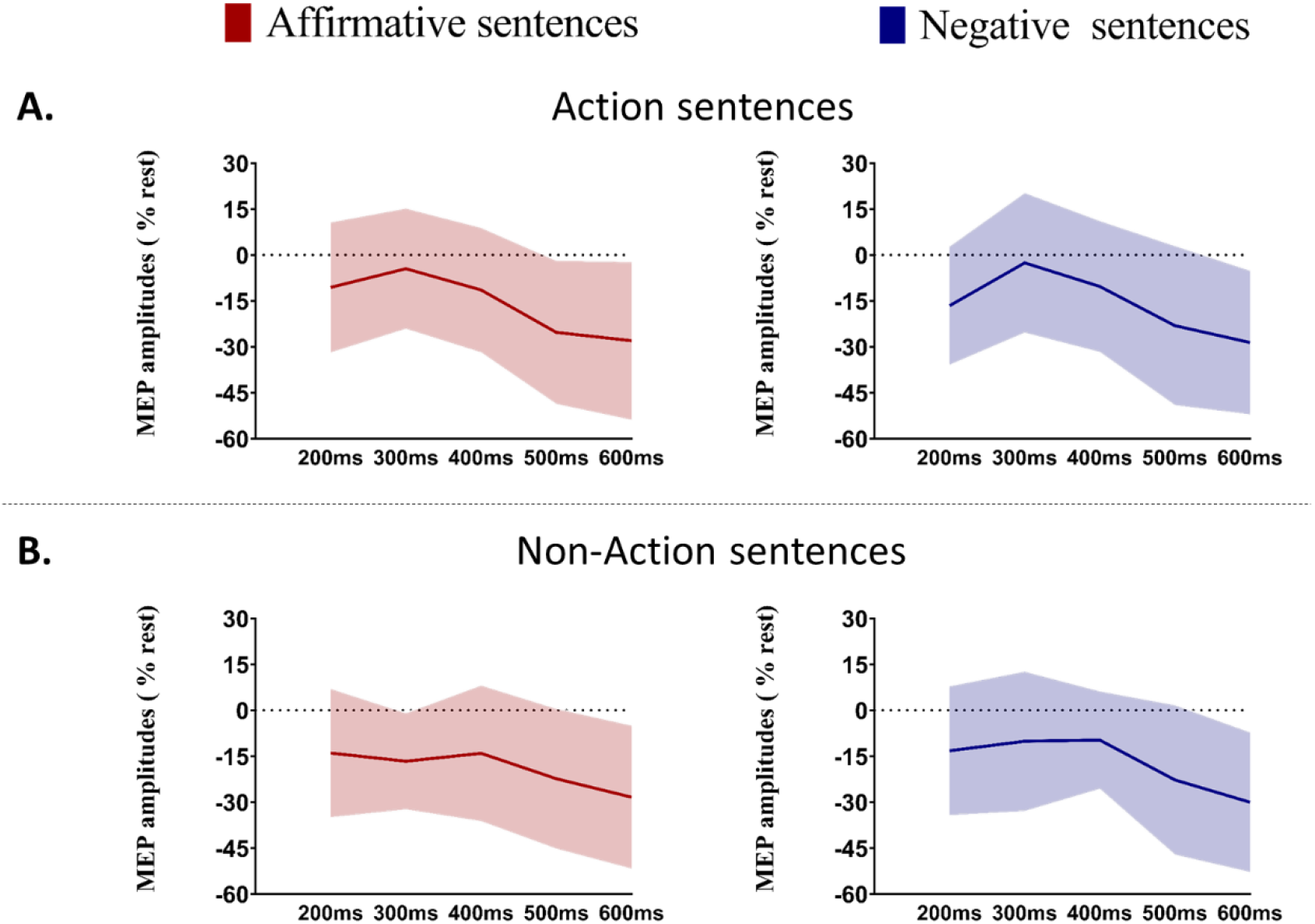
Corticospinal excitability for action and non-action sentences in affirmative and negative forms. Lines represent mean amplitude and shaded areas denote the standard deviation.

Moreover, we qualitatively observed that all conditions exhibited a MEP amplitude peak and crucially that the latencies preceding and/or following this peak yielded a decrease in corticospinal excitability. This suggests that the involvement of the motor system is punctual (See Figure 3).

**Figure 3:**
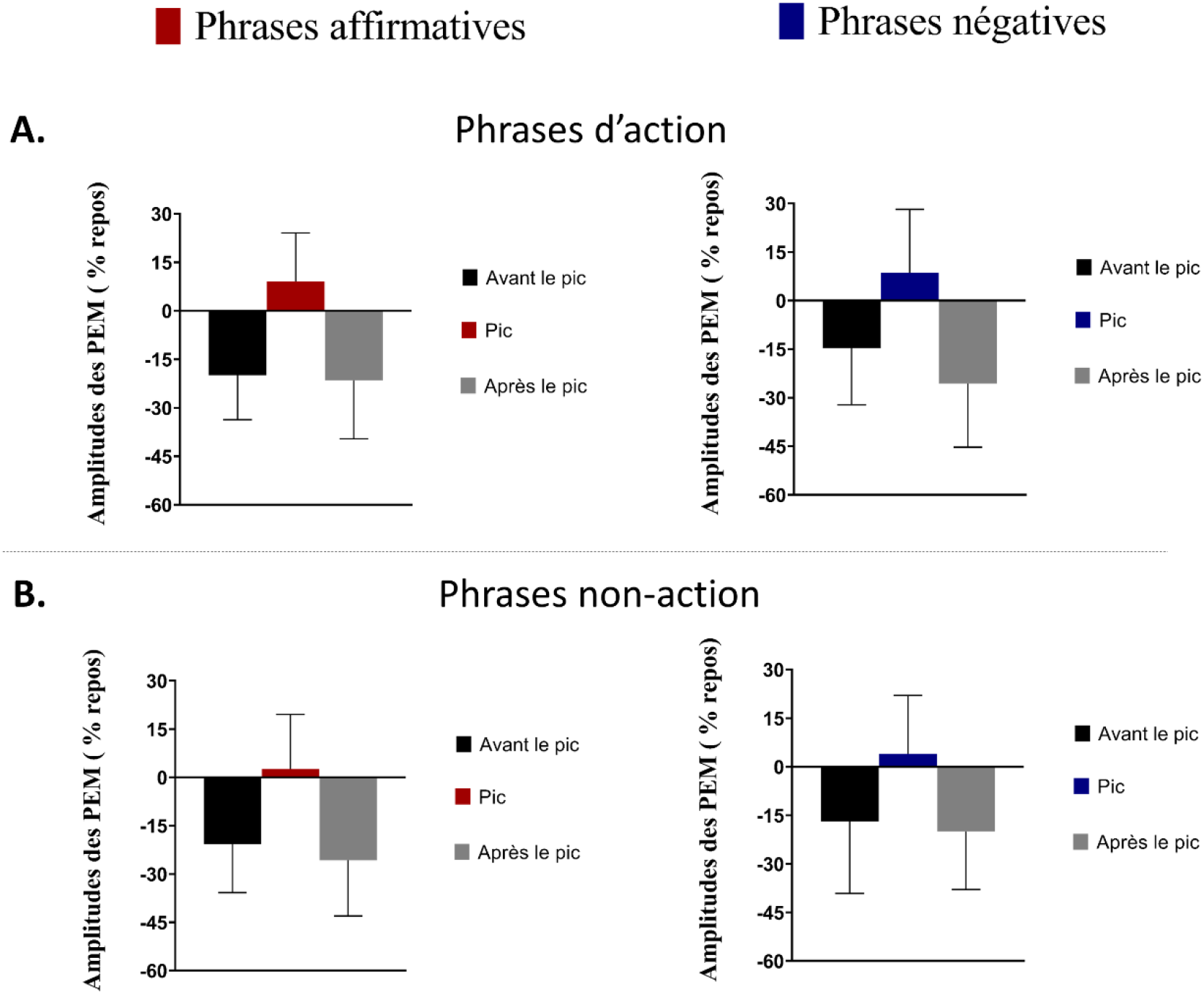
Temporal dynamics of the corticospinal excitability surrounding the peak MEP amplitude. Bar plots denote the mean and vertical bars represent the standard deviation.

Using individual peaks (i.e., only the one of the three latencies yielding the highest MEP for a condition), normalized MEPs for affirmative and negative action sentences were different from rest (affirmative: p=0.007; t=2.938; negative sentences: p=0.044; t=2.121), but non-action sentences were not (affirmative: p=0.453; t=0.762; negative: p=0.290; t=1.081). Moreover, a main effect of Action was observed (F_1, 23_=4.865, p=0.037, ηp^2^=0.174), with larger MEP amplitude for Action (8.77 ±14.87%) than Non-action sentences (3.31 ±15.75%, Cohen’s d=0.36). However, we did not observe any Polarity effect (F_1, 23_=0,031, p=0.860, ηp^2^=0.001) nor interaction between Action and Polarity (F_1, 23_=0,134, p=0.717, ηp^2^=0.005) (Figure 4).

**Figure 4:**
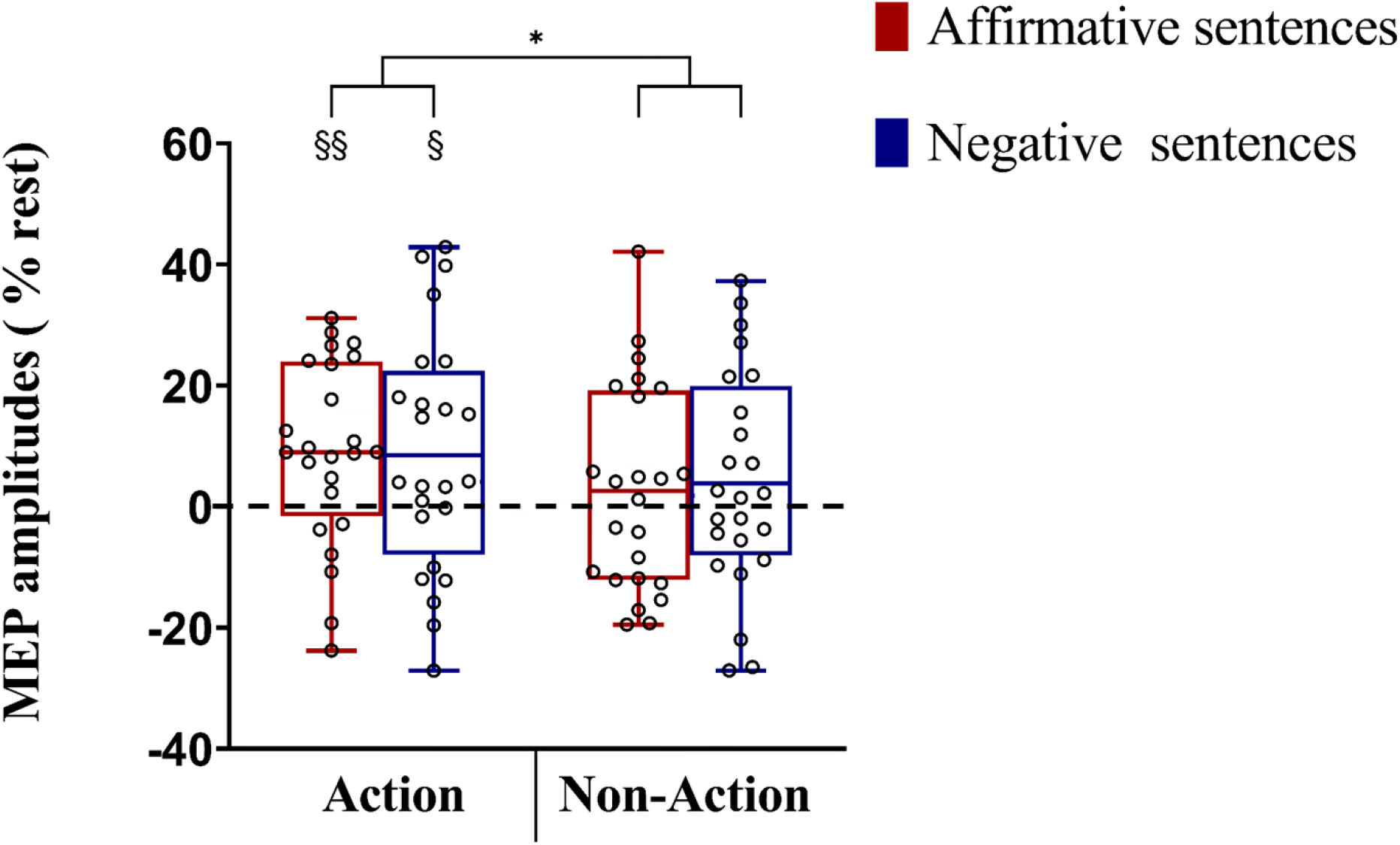
Box plots represent normalized MEPs (% rest). Circles represent individual data. The rmANOVA yielded a main effect of action. *=p<0.05. §= difference from rest.

These findings demonstrate greater corticospinal excitability for action sentences compared to non-action sentences and rest. Nevertheless, this state of excitability does not seem to be polarity-dependent, meaning that action verbs in the affirmative and negative form do not induce different modulations of the corticospinal pathway at these latencies.

### Experiment 2

In this second experiment, we only presented affirmative and negative action sentences (no non-action sentences) at short and long latencies.

#### Corticospinal excitability

Due to inter-subject variability of latencies, we selected individual peak MEP increase, as in Experiment 1. We found that corticospinal excitability increased for all conditions in comparison to rest (all p’s <0.05). However, we did not find any significant main effect of Polarity (F_1,18_=0.146, p=0.706, ηp^2^=0.008), Latency (F_1,18_=1.867, p=0.188, ηp^2^=0.093), nor Latency by Polarity interaction. (F_1,18_=0.006, p=0.936, ηp^2^=0.000) (See Figure 5).

**Figure 5:**
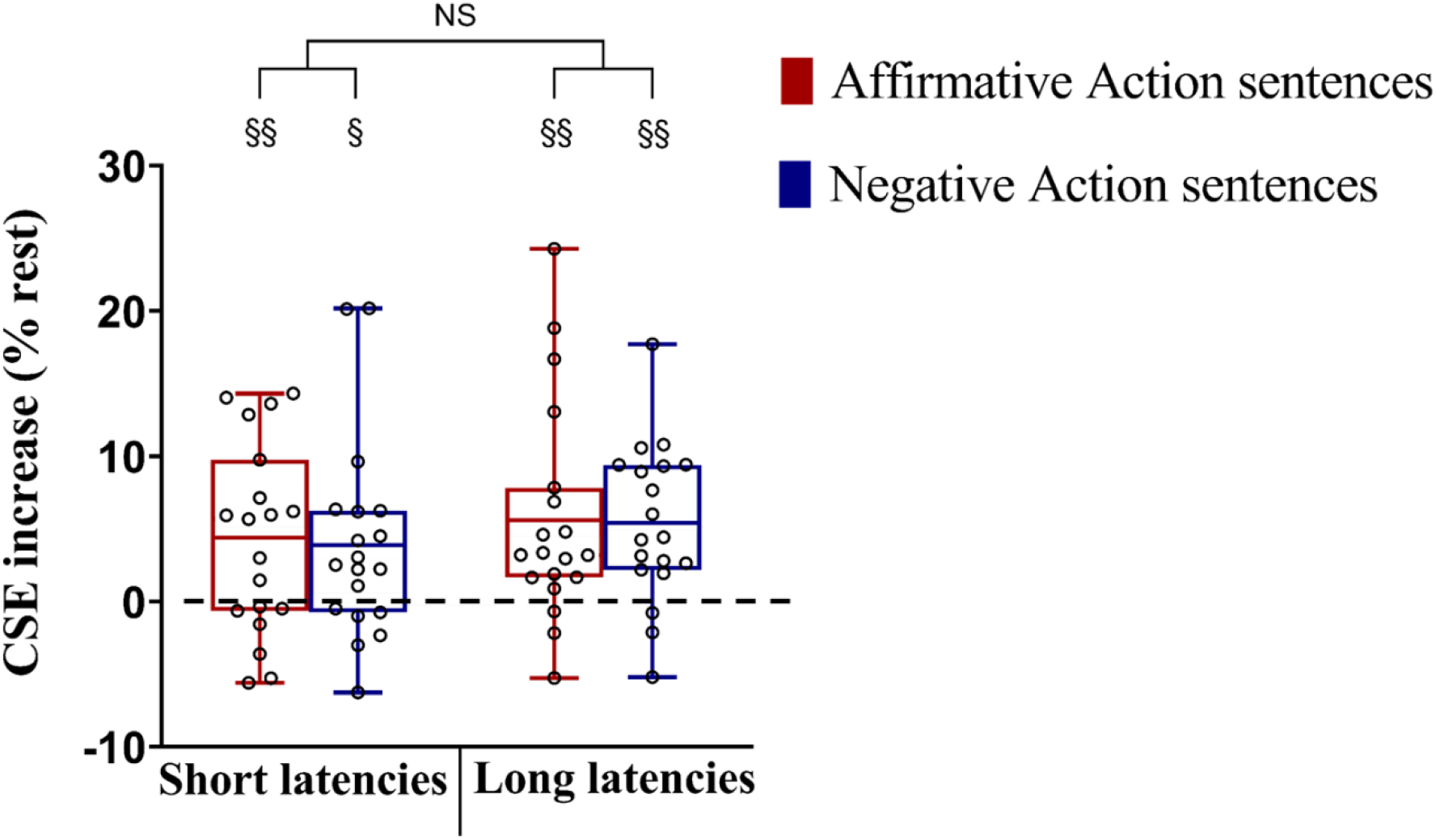
Corticospinal excitability (CSE) at short and long latencies in Experiment 2. Box plots represent corticospinal excitability (percentage rest). Circles represent individual data.

These findings show that affirmative and negative action sentences led to a greater peak of corticospinal excitability in comparison to rest, but this increase did not differ from early to late latencies. In summary, both affirmative and negative action sentences engage the motor system at both latencies tested in the current study.

#### Short-interval intracortical inhibition (SICI)

First, we found a higher TS intensity to reach the target MEP amplitude for paired-pulse in comparison to single pulse stimulation at rest, reflecting inhibition within M1 (25.64%; p<0.001; t=10.243). Next, using the same peak latency from the corticospinal excitability results, we found a larger inhibition at the long latency than at rest (for affirmative sentences, p=0.005; t=3.160 and for negative sentences, p=0.002; t=3.637), whereas the short latency yielded no supplementary inhibition (for affirmative sentences, p=0.218; t=1.276 and for negative sentences, p=0.234; t=1.231). Moreover, we observed a main effect of Latency (F_1,18_=12.968, p=0.002, ηp^2^=0.418) with greater inhibition at long (12.057 ±15.723%) than short latencies (4.455 ±15.542%, Cohen’s d=0.50). However, we did not observe any significant main effect of Polarity (F_1,18_=0.499, p=0.488, ηp^2^=0.027), nor Latency by Polarity interaction. (F_1,18_=0.756, p=0.395, ηp^2^=0.040) (Figure 6).

**Figure 6:**
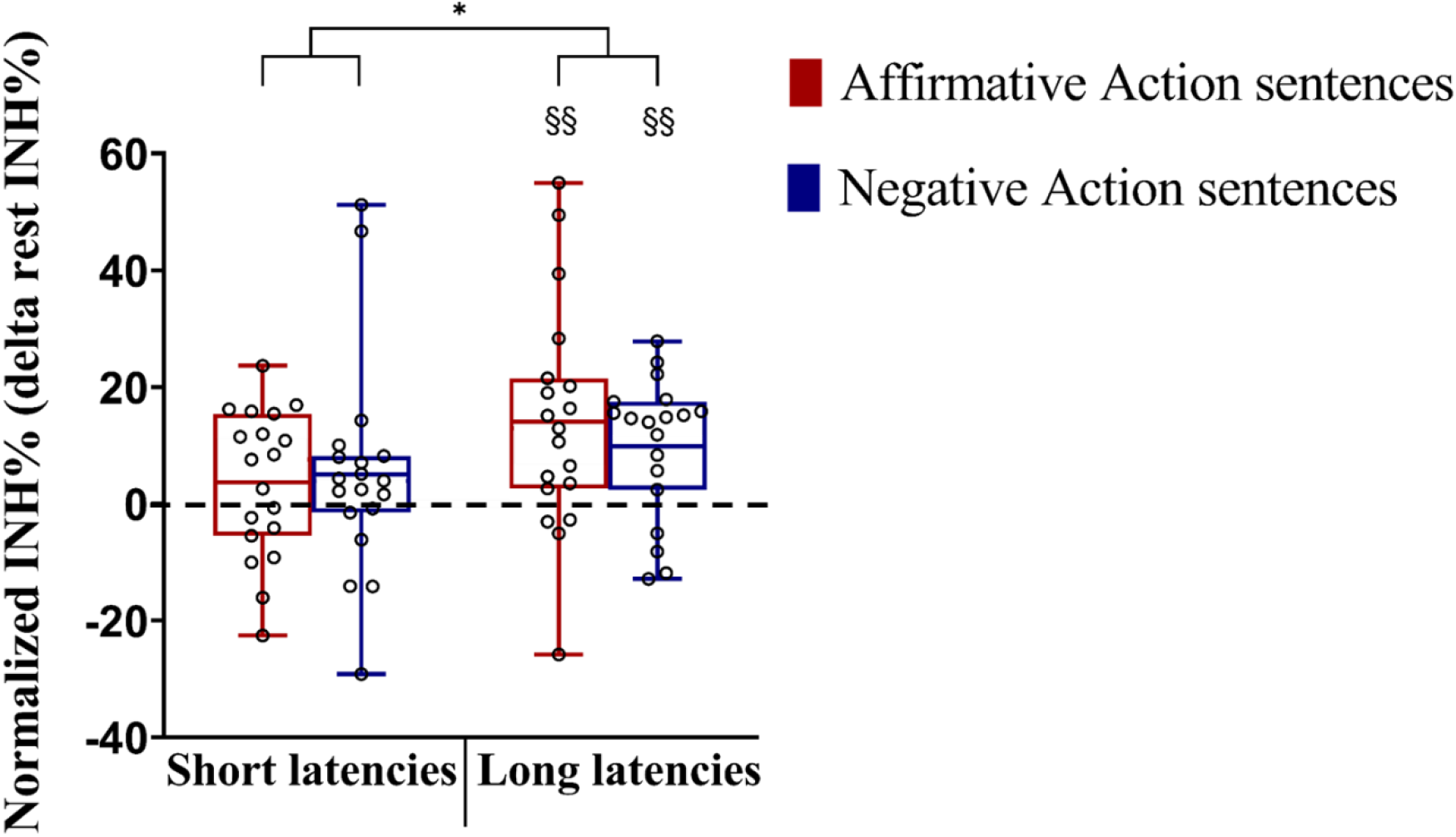
SICI ratio for affirmative and negative action sentences at short and long latencies in Experiment 2. Box plots represent intracortical inhibition (delta to rest intracortical ratio). Circles represent individual data.

Taken together, these results suggest that negative action sentences do not immediately lead to greater intracortical inhibition in comparison to affirmative action sentences within M1. At later latencies, inhibition is augmented for both negated and affirmed actions.

## Discussion

The present study describes two original findings regarding the comprehension of negated actions. The first result of interest is the increase of corticospinal excitability for both affirmative and negative action sentences at short and long latencies following verb presentation. The second result is the increase of SICI within M1 at longer latencies, regardless of sentence polarity.

### Increased corticospinal excitability at early latencies for both affirmative and negative actions

The increase of corticospinal excitability observed in this study suggests that even negative action sentences may be simulated in a way that engages the motor system. This finding supports the initial step of the two-step theory of negated language processing (Beltrán et al. 2021; Kaup et al. 2005, 2006; Kaup, Yaxley, et al. 2007; Lüdtke et al. 2008; Anderson et al. 2010; Scappini and Delfitto 2015). The idea in this two-step theory is that negated actions are mentally simulated in a first step, and afterward suppressed in a second step. At the same time, our failure to observe a later difference between negated and affirmed actions contradicts recent research showing a decrease of corticospinal excitability after reading negative compared to affirmative action verbs (Papeo et al. 2016) or adverbs (Papitto et al. 2021). There are various potential reasons for this discrepancy. First, there are differences in the linguistic stimuli and the method of presentation between these studies. In the current study, we employed complex sentences presented in two clauses, whereas Papeo et al. (2016) and Papitto et al. (2021) used minimal context (short verb phrases, e.g. “now I write”) and a rapid serial visual presentation, respectively. The richer context and more natural reading presentation may lead to a stronger motor simulation. Secondly, measurement timing is a factor of significant interest in action reading literature (Buccino et al. 2005; Borreggine and Kaschak 2006; Kaschak and Borreggine 2008; Papeo et al. 2009), which could also explain discrepancies. In our study, we probed a wider variety of TMS stimulation latencies, and addressed individual variability in activation of the action representation by performing a novel peak analysis. If we hadn’t considered individual latencies of corticospinal excitability increase, we may have missed the motor system involvement during the processing of negative action verbs. The combination of these factors may explain that we observed an increase of corticospinal excitability after reading a negative sentence.

### A late peak in corticospinal excitability for both affirmative and negative actions

The increase of corticospinal excitability for both negative and affirmative action sentences at long latencies was unexpected. While the early increase of corticospinal excitability can be explained by a facilitation within M1, arising from the motor representations evoked during the reading of action sentences, the origins of the later peak are more difficult to pinpoint. The later increase in corticospinal excitability could arise from a second, later stage of processing of the described action (or a re-processing) within the motor system. On the other hand, it could also reflect inputs from other facilitatory processes originating in regions external to M1. This latter facilitation may be transmitted from regions such as premotor or parietal areas, or even from subcortical or spinal levels of processing (Reis et al. 2008). If this second peak indeed arises from regions external to the motor system, the delay could be driven by the time needed for initial processing within these regions, as well as the time to transmit signals through direct or indirect pathways between these regions and M1 or the corticospinal pathway.

### Increased inhibition for both affirmative and negative actions at long latencies

We observed an increase in intracortical inhibition within M1 for negated sentences at longer latencies (compared to rest and shorter latencies), which would seem to support the two-step processing theory during negative reading (Beltrán et al. 2021; Kaup et al. 2005, 2006; Kaup, Yaxley, et al. 2007; Lüdtke et al. 2008; Anderson et al. 2010; Scappini and Delfitto 2015). However, the fact that the same inhibitory pattern was observed for affirmative action sentences was unexpected. One hypothesis for these inhibitory mechanisms during the reading of action sentences could be to limit motor system activation in order to impede unwanted movements. This hypothesis has been put forth for motor imagery (Jeannerod and Decety 1995; Jeannerod 2001), yielding similar results for corticospinal excitability and SICI (Neige et al. 2020).

The increase of corticospinal excitability at long delays, combined with the simultaneous increase of SICI for both negated and affirmed sentences might be explained by the fact that corticospinal excitability comprises inhibitory and excitatory processes inside the corticospinal tract including both cortical and spinal contributions (Neige et al. 2020). As discussed above, corticospinal facilitation at long latencies could potentially originate outside M1 and exert its influence through direct or indirect pathways (Reis et al., 2008). By contrast, the modulation of SICI observed in our study reflects cortical inhibitory interneurons within M1. Given that inhibition within M1 is greater at long than short latencies, it is possible that late inhibition is required to counter increases in corticospinal excitability arriving from other premotor, parietal or subcortical regions.

The idea of two-step processing of negated actions specifies that negated actions are mentally simulated and afterward suppressed by recruiting general action inhibitory mechanisms (e.g., Beltrán et al. 2021). In our study, while the predicted increase in cortical excitability for negated sentences was observed at the earlier latencies, no difference in cortical excitability nor inhibition between negated and affirmed sentences was observed at longer latencies, at least not within M1 and along the corticospinal pathway. Nevertheless, we must be mindful that the observed lack of difference in inhibition refers specifically to the inhibitory mechanism within M1. Negated concepts may well be inhibited in other systems in the brain such as the dorsolateral prefrontal cortex (dlPFC), the pre-supplementary motor area (pre-SMA), the inferior parietal or the inferior frontal gyrus. Recent evidence from EEG studies supports distinct inhibitory mechanisms in the inferior frontal gyrus/pre-SMA during negated and affirmative action reading (De Vega et al. 2016; Beltrán et al. 2018, 2019, 2021; Liu et al. 2020; Vitale et al. 2021). These findings permit the speculation that various brain regions and systems contribute differentially to the two stages of processing for negated actions. It is possible that some regions, such as M1, would contribute to the initial processing step; engaging a representation or simulation of the negated action. This is essential for the ability to understand what has been negated, and can be likened to a sort of motoric lexical access of the negated concept. It is possible that this information is then relayed to other regions or multi-sensory integration areas where this concept can be inhibited to capture the intended negation.

### Perspectives and limitations

There are several limitations of the present research. First, there were a limited number of experimental trials per participant. As it remains impossible to know exactly when the motor representation will occur and reach its maximum for each subject and each trial, we had to probe several latencies on separate trials, and select the peak afterwards, discarding the trials that missed the motor facilitation. This could be considered a strength of the current study in that individual differences in processing time are taken into account, but it can also be a limitation in that we reduce the number of experimental trials. We attempted to overcome this limitation in Experiment 2 by calibrating each subject’s individual processing time prior to the test session, and then using that latency during the experimental trials. This procedure was used only for the short latencies, as we considered the facilitation and inhibition at longer latencies to be more susceptible to variations, and thus less stable. We are looking into continuous measures, such as EEG, to further investigate these issues.

A second limitation is the restriction of our measurements to the motor system. Given the study’s focus on action language processing and our interest in the motor simulation, this is a deliberate and suitable choice. However, our results raise questions that implicate regions outside of M1, and further research will indeed be necessary to address the possible inhibition of the negated actions at other brain sites. It would also be interesting to determine whether the increase in corticospinal excitability in the long latencies originates from the motor simulation in M1 or from other brain regions. For example, double-site TMS might be useful in investigating the interaction between M1 and the inferior parietal lobule, which plays a role in comprehension as well as action inhibition.

### Conclusions

In conclusion, the present study provides relevant information about the processing of negated actions within M1 and along the corticospinal track. The augmentation in corticospinal excitability observed at early latencies suggests that even negative action sentences may be simulated in a way that engages the motor system. This finding is consistent with the initial step of the two-step theory of negated language processing, specifying that negated actions are mentally simulated in a first step. The second step of this theory postulated the suppression of this simulation. At later latencies, we indeed observed increased SICI, but this was the case for both negated and affirmed sentences. Therefore, it is more likely that this inhibition process serves to counter motor facilitation and impede movement during motor cortex activation, as postulated in motor imagery, rather than to annul the negated concept. It remains possible that a distributed multimodal network is required to capture the semantic nature of a negated concept, such that the motor system helps to represent an action to be negated, and a subsequent inhibitory process comes to play on this representation in a separate brain region. While the current study lends support for the motor system’s role in the initial stage of the two-step theory of negation processing, further research is clearly needed to better specify the second step and the contribution of other brain regions to this process.

## Supporting information

Supplemental section

## Author Contributions

Experiment design: WD, CML, FL; Data collection: WD, LL; Statistical analysis: WD, CML, FL; Manuscript preparation: WD, CP, CML, FL

## Declaration of competing interest

None declared.

## Notes

### Competing Interest Statement

The authors have declared no competing interest.

