## Supplemental section for "Reading negative action verbs: one or two-step processing within the primary motor cortex?"

### Supplementary section

#### Method

*Table 1: Example of lexical stimuli used in experiment.*

| Action | | Non-action | |
|---|---|---|---|
| **Affirmative** | **Negative** | **Affirmative** | **Negation** |
| La pomme de terre est cuite, je l'épluche | La pomme de terre est cuite, je ne l'épluche pas | La pomme de terre est cuite, je la laisse | La pomme de terre est cuite, je ne la laisse pas |
| *The potato is cooked, I peel it* | *The potato is cooked, I don't peel it* | *The potato is cooked, I leave it* | *The potato is cooked, I don't leave it* |
| Le chat ronronne, je le caresse | Le chat ronronne, je ne le caresse pas | Le chat ronronne, je l'imite | Le chat ronronne, je ne l'imite pas |
| *The cat purrs, I caress it* | *The cat purrs, I don't caress it* | *The cat purrs, I imitate it* | *The cat purrs, I don't imitate it* |
| Le son de la guitare est mauvais, je l'accorde | Le son de la guitare est mauvais, je ne l'accorde pas | Le son de la guitare est mauvais, je l'écoute | Le son de la guitare est mauvais, je ne l'écoute pas |
| *The sound of the guitar is bad, I tune it.* | *The sound of the guitar is bad, I don't tune it.* | *The sound of the guitar is bad, I listen it.* | *The sound of the guitar is bad, I don't listen it.* |
| Je réfléchis à la question, j'écris | Je réfléchis à la question, je n'écris pas | Je réfléchis à la question, j'hésite | Je réfléchis à la question, je n'hésite pas |
| *I think about the question, I write* | *I think about the question, I don't write* | *I think about the question, I hesitate* | *I think about the question, I don't hesitate* |
| Mon immeuble a un code d'entrée, je le tape | Mon immeuble a un code d'entrée, je ne le tape pas | Mon immeuble a un code d'entrée, je le retiens | Mon immeuble a un code d'entrée, je ne le retiens pas |
| My building has an entry code, I type it | My building has an entry code, I don't type it | My building has an entry code, I remember it. | My building has an entry code, I don't remember it. |

*Table 2: T-test results concerning the linguistic and psycholinguistic characteristics of stimuli.*

| Factors | Means<br>Action sentences | Means<br>Non- Action sentences | T-tests<br>(corrected p-value) |
|---|---|---|---|
| **Written frequency** | 9.57±13.93 | 16.94±35.09 | 0.30 (t=1.88) |
| **Number of syllables** | 1.66±0.54 | 1.82±0.56 | 0.45 (t=1.70) |
| **Number of characters** | 6.36±1.39 | 6.40±1.51 | 1 (t=-0.05) |
| **Spelling neighbors** | 5.26±4.63 | 4.75±5.01 | 1 (t=-0.62) |
| **Semantic analyse** | 0.32±0.29 | 0.31±0.28 | 1 (t=-0.25) |

*Table 3: Distribution of the MEP peak latency.*

| latency | Affirmative Action sentences | Negative Action sentences | Affirmative Non-Action sentences | Negative Non-Action sentences |
|---|---|---|---|---|
| **200ms** | 6 | 3 | 5 | 9 |
| **300ms** | 9 | 13 | 7 | 6 |
| **400ms** | 5 | 3 | 9 | 4 |
| **500ms** | 2 | 3 | 1 | 4 |
| **600ms** | 2 | 2 | 2 | 1 |

#### Results EMGrms

##### Experiment 1

We analyzed the EMGrms before TMS artifacts to ensure MEP modulation was not influenced by muscle contractions. We did not observe any difference between our conditions (Friedman ANOVA: $p=0.283$, $r=0.011$). This result demonstrates that MEP modulations were not influenced by muscular pre-activity.

*Table 1: EMGrms activity (mean ±SD) in µV recorded for the first dorsal interosseous (FDI) before the single TMS artifact for each condition (window of 100ms prior the artifact).*

| | | **Affirmative form** | | **Negative form** | |
|---|---|---|---|---|---|
| **Conditions** | Rest | Non-action | Action | Non-action | Action |
| **Mean ±SD (µV)** | 1.56 ± 0.68 | 1.50 ± 0.72 | 1.53 ± 0.77 | 1.56 ± 0.79 | 1.61 ± 0.89 |

##### Experiment 2

We analyzed the EMGrms before TMS artifacts to ensure MEP modulation was not influenced by muscle contractions. We did not observe any difference between our conditions (Friedman ANOVA: For corticospinal excitability $p=0.367$, $r=0.004$; for SICI $p=0.617$, $r=-0.018$). This result demonstrates that MEP modulations were not influenced by muscular pre-activity.

*Table 2: EMGrms activity (mean ±SD) in µV recorded for the first dorsal interosseous (FDI) before the single and paired TMS artifact for each condition (window of 100ms prior the artifact).*

| | **Corticospinal excitability** | | | | |
|---|---|---|---|---|---|
| | | **Affirmative form** | | **Negative form** | |
| **Conditions** | Rest | Short latency | Long latency | Short latency | Long latency |
| **Mean ±SD (µV)** | 0.958 ± 0.356 | 1.067 ± 0.589 | 1.254 ± 0.775 | 1.071 ± 0.804 | 1.099 ± 0.788 |
| | **Short IntraCortical Inhibition (SICI)** | | | | |
| | | **Affirmative form** | | **Negative form** | |
| **Conditions** | Rest | Short latency | Long latency | Short latency | Long latency |
| **Mean ±SD (µV)** | 1.094 ± 0.559 | 1.126 ± 0.581 | 0.991 ± 0.443 | 0.937 ± 0.274 | 0.902 ± 0.283 |
